# Enterovirus-driven interferon signaling induces epithelial TG2 via JAK-STAT: Implications for the onset of celiac disease

**DOI:** 10.64898/2026.05.26.727875

**Authors:** Hana Hien Le, Veera Räkköläinen, Roselia Davidsson, Valeriia Dotsenko, Laura Martin Diaz, Amirbabak Sioofy Khojine, Anniina Virtanen, Jutta E. Laiho, Chaitan Khosla, Olli Silvennoinen, Heikki Hyöty, Keijo Viiri, the HEDIMED Investigator group

**Affiliations:** Celiac Disease Research Center, Faculty of Medicine and Health Technology, Tampere. University, and Tampere University Hospital, Tampere, Finland; Faculty of Medicine and Health Technology, Tampere University, Tampere, Finland; Departments of Chemistry and Chemical Engineering and Sarafan ChEM-H, Stanford University, Stanford CA 94305.; Fimlab laboratories, Tampere, Finland; Institute of Biotechnology, HiLIFE Helsinki Institute of Life Science, University of Helsinki; Faculty of Biochemistry and Molecular Medicine, University of Oulu, Oulu, Finland

**Author notes:** Correspondence. Address correspondence to: Keijo Viiri, PhD, Celiac Disease Research Center, Faculty of Medicine and Health Technology, Tampere University Hospital, Tampere University, Arvo Ylpön Katu 34, Tampere, FIN-33520, Finland. List of HEDIMED investigators is appended as a separate document.

## Abstract

**Background & Aims:** Celiac disease (CeD) is an autoimmune disorder triggered by dietary gluten in genetically predisposed individuals, but environmental factors contributing to disease onset remain incompletely defined. Epidemiological studies implicate enterovirus infections as potential triggers. Here, we investigated the epithelial-intrinsic mechanisms by which coxsackievirus B1 (CVB1) infection may prime the intestine for CeD.

**Methods:** Human intestinal organoids were infected with CVB1 and analyzed using single-cell RNA sequencing to resolve lineage-specific responses. Interferon signaling and transglutaminase 2 (TG2) regulation were interrogated using type I interferon stimulation and pharmacologic JAK inhibition.

**Results:** CVB1 infection induced a robust epithelial antiviral program dominated by type I interferon signaling. This response was accompanied by marked upregulation of TG2 expression and enzymatic activity. Single-cell analysis localized TG2 induction to immature goblet-lineage cells, which exhibited strong interferon-stimulated gene activation and epithelial stress signatures. Mechanistically, IFN-α/β stimulation was sufficient to induce TG2 via JAK–STAT signaling, while JAK inhibition effectively suppressed both TG2 expression and activity. In parallel, CVB1 infection triggered coordinated mucin remodeling, including induction of MUC5AC, indicating interferon-linked epithelial reprogramming. Notably, these effects occurred independently of immune cell involvement, highlighting a cell-intrinsic pathway.

**Conclusion:** Our findings identify a direct epithelial mechanism linking enterovirus infection to TG2 activation via interferon-driven JAK–STAT signaling. This pathway provides a mechanistic bridge between viral infection and gluten peptide modification, a critical step in the onset of CeD. The reversibility of TG2 induction by JAK inhibition suggests a potential strategy to prevent virus-mediated priming of celiac disease.

## INTRODUCTION

Celiac disease (CeD) is a chronic immune-mediated enteropathy affecting up to 2% of the population and is triggered by dietary gluten in genetically predisposed individuals carrying HLA-DQ2 or HLA-DQ8 haplotypes.^1^ Although these genetic factors are necessary, they are not sufficient to drive disease onset, highlighting the importance of environmental triggers.^2^ Currently, the only effective treatment is lifelong adherence to a gluten-free diet, underscoring the need to better understand the mechanisms initiating loss of tolerance to gluten.

A central event in CeD pathogenesis is the modification of gluten peptides by the enzyme transglutaminase 2 (TG2). TG2 deamidates specific glutamine residues in gluten, generating negatively charged peptides with increased affinity for HLA-DQ2/DQ8, thereby promoting activation of pathogenic CD4+ T cells and mucosal inflammation.^3–7^ The critical role of TG2 in disease has been demonstrated therapeutically, as pharmacologic inhibition of TG2 attenuates gluten-induced intestinal injury^8,9^ and also restore gluten-induced systemic changes in the plasma.^10^ Despite its importance, the upstream mechanisms regulating TG2 expression and activity in early disease initiation remain incompletely defined.

Emerging evidence suggests that environmental factors, particularly viral infections, may contribute to the breakdown of oral tolerance to gluten. Prospective cohort studies have implicated enteroviruses as potential triggers of CeD, and viral signals have been detected in the intestinal mucosa of affected individuals.^11–14^ However, the molecular mechanisms linking viral infection to CeD pathogenesis remain unclear. In particular, whether enterovirus-induced epithelial responses can directly regulate TG2 and thereby promote disease-relevant biochemical changes has not been resolved.

The intestinal epithelium forms the first line of defense against luminal pathogens and actively coordinates innate immune responses. Upon viral infection, epithelial cells rapidly produce type I and III interferons, activating antiviral programs through the JAK–STAT signaling pathway.^15^ While interferon-γ has been shown to induce TG2 expression,^9,16^ it is unknown whether epithelial-intrinsic antiviral interferon responses triggered during viral infection similarly drive TG2 activation. Defining such a mechanism would provide a direct link between viral exposure and the key enzymatic step underlying gluten immunogenicity.

Here, we hypothesized that enterovirus infection induces epithelial TG2 activity through interferon-driven JAK–STAT signaling. Whereas upregulation of TG2 expression has been observed in multiple studies involving active CeD patients^9,17,18^, direct measurement of TG2 activity in vivo is technically challenging due to the existence of elaborate allosteric regulatory mechanisms in this mammalian enzyme. Using human intestinal organoids and single-cell transcriptomics, we investigated how coxsackievirus B1 (CVB1) infection reshapes epithelial cell states and signaling networks. Our findings reveal a lineage-specific, epithelial-intrinsic pathway in which antiviral interferon responses drive TG2 expression and activity, providing a mechanistic bridge between enterovirus infection and early events in CeD pathogenesis.

## MATERIALS AND METHODS

### Human intestinal organoid culture

Samples were collected from duodenal biopsies of non-celiac patients from the Tampere University Hospital. Isolation and culture of epithelial organoids were performed as described previously.^19^ Briefly, isolated crypts were mixed with Matrigel (Corning Inc., New York, USA) and cultured in ADF medium which included DMEM/F12 medium (Thermo Fisher Scientific, Waltham, MA, USA), 100 U/ml penicillin and 100 μg/ml streptomycin (Thermo Fisher Scientific), 10 mM HEPES (Thermo Fisher Scientific), and 2 mM GlutaMax (Thermo Fisher Scientific), supplemented with 10% v/v WNT3A-conditioned medium, 500 ng/ml human R-spondin (R&D Systems, Inc., MN, USA), 0.2 μM LDN (Sigma-Aldrich, Saint Louis, MO, USA), 1,25 mM N-acetylsysteine (Sigma-Aldrich), 10 mM Nicotinamide (Sigma-Aldrich), 3 μM SB202190 (Sigma-Aldrich), 1x B27 (Sigma-Aldrich), 50 ng/ml human EGF (Thermo Fisher Scientific), 500 mN A-83-01 (Sigma-Aldrich), and 1 μM Prostaglandin E2 (Sigma-Aldrich). The organoids were cultured with a split ratio of 1:2 every 7 days. For epithelial cell differentiation, the organoids were grown in ADF medium with the supplements of 2% v/v B27, 1.25 mM N-acetylcysteine, 0.1 μM LDN, 250 ng/ml human R-spondin, and 50 ng/ml human EGF for 48 hrs incubation.

### Organoid infection

Organoids were moved to the differentiation medium 2 days prior to virus infection. At the time of infection, the medium was removed, and the organoids were resuspended in cold Phosphate buffered saline (PBS) to remove any excess of Matrigel. To enhance infection efficiency, the organoids were disrupted in Cell Recovery Solution (Corning, NY, USA) for 30 min at 4°C with constant agitation to dissolve the Matrigel. Mock-infected cells were incubated with media whereas infected samples were incubated with supernatant containing viruses with an MOI of 6 for acute infection. Infections were proceeded for 1 h at room temperature (RT). Cells then spun down at 400 g for 5 min at RT to remove the viruses and washed 3 times with PBS. The cells were then embedded in Matrigel in differentiation media for the course of the experiment for 24 hrs.

### Organoid treatments

Non-infected organoids were maintained in ADF medium for 2-3 days to allow for expansion. All inhibitor treatments were performed in the presence of 5 ng/mL IFN-α (Peprotech, Rocky Hill, NJ, USA) or IFN-β (Peprotech), together with individual JAK inhibitors, momelotinib (Selleck Chemicals, Houston, TX, USA), tofacitinib (Selleck Chemicals), deucravacitinib (MedChemExpress, Monmouth Junction, NJ, USA), ritlecitnib (Sigma-Aldrich) and VVD-118313 (Vividion, San Diego, CA, USA), at a final concentration of 500 nM.

### Western blot

Organoids were treated with JAK inhibitors for 1h prior to IFN stimulation for 15 minutes. Organoids were harvested in Cell Recovery solution (Corning), washed with 1x PBS and resuspended in RIPA-buffer containing protease and phosphatase inhibitors. Lysates were clarified by centrifugation and stores at –20 °C for further analysis.

Samples were mixed with 4xSDS-PAGE buffer, boiled at 95 °C for 5 minutes, separated on Bio-Rad TGX 7.5% mini gels and transferred to nitrocellulose membranes using the Trans-Blot Turbo (Bio-Rad) system. Membranes were blocked with EveryBlot (Bio-Rad) buffer and probed with mouse anti-pSTAT1 (clone ST1P-11A5, Thermo Fisher Scientific), rabbit anti-STAT1 (clone 9172, CST) and mouse anti-β-tubulin (clone sc-55529, Santa Cruz) (1:1000), followed by DyLight 800 anti-mouse and DyLight 680 anti-rabbit secondary antibodies (Thermo Fisher Scientific). Blots were imaged on the Odyssey CLx system, and band intensities quantified using Image Studio software.

### Confocal immunofluorescent microscopy

After 24 h JAK inhibitor treatment TG2 enzymatic activity was visualized using the fluorescent probe HB-2-30^20^ (Biotechne Tocris, Cat.No.8831, USA) according to the manufacturer’s protocol with Nikon A1R+ Laser Scanning confocal microscope.

### RNA isolation

Cultured medium and organoids were harvested 24 hours after infection or JAK inhibitor treatment and then mRNA was isolated using TRIzol™ Reagent (Thermo Fisher Scientific) following the manufacturer’s protocol. Briefly, samples were mixed with TRIzol™ Reagent by pipetting several times to homogenize and then incubated for 5 min to allow complete dissociation of the nucleoprotein complex. Chloroform (Thermo Fisher Scientific) was added to the samples for mRNA collection. mRNA was washed twice with 70% ethanol and stored at -80°C before further processing.

### Viral RNA extraction and quantification

Purified CVBs supernatant was aliquoted and serially diluted in sterile PBS to generate a dilution series for downstream RNA quantification. The viral RNA was extracted from each dilution with TRIzol reagent (Thermo Fisher Scientific) according to the manufacturer’s protocol. Briefly, 250 μL of virus supernatants were mixed with 500 μL of Trizol reagent, then incubated at RT for 15 min, and followed by phase separation using chloroform. RNA was precipitated with isopropanol, washed with 75% ethanol, and resuspended in RNase-free water. RNA concentrations were measured using a Nanodrop spectrophotometer (Thermo Fisher Scientific, USA). The samples were stored at -80°C for further processing.

### RT-qPCR

cDNA was made using iSCRIPT reverse transcriptase (Bio-Rad) from 1 μg of total RNA per sample following the manufacturer’s instructions. qPT-PCR was performed using SsoFast™ EvaGreen® Supermix (Bio-Rad) with GAPDH used as normalizing genes. Virus copies were calculated based on the standard curve as described previously.^21^ The specific primers ordered from Sigma-Aldrich are as follows:

CVB1 forward, 5’-CCCTGAATGCGGCTAATCC-3’ and reverse, 5’-ATTGTCACCATAAGCAGCCA-3’;

GAPDH forward, 5’-TCCATGACAACTTTGGTATCGTGG-3’, and reverse, 5’-GACGCCTGCTTCACCACCTTCT-3’;

IFN-β forward, 5’-GCCGCATTGACCATCTAT-3’, and reverse, 5’-GTCTCATTCCAGCCAGTG-3’;

IFN2/3 forward, 5’-GCCACATAGCCCAGTTCAAG-3’, and reverse, 5’-TGGGAGAGGATATGGTGCAG-3’;

TG2 forward, 5’-AGGAGATGCCACCTGTTGAG -3’, and reverse, 5’-TCACAGAGCAGCCACATCAA-3’;

STAT1 forward, 5’-CGCAGAAAAGTTTCATTTGCTGT-3’, and reverse, 5’-CCACTGAGACATCCTGCCAC-3’;

GBP1 forward, 5’-CAAGGGAACAGCCTGGACAT-3’, and reverse, 5’-GCCCAGAGAGAAGCCCTTTTT-3’;

and IRF1 forward, 5’-GAGGAGGTGAAAGACCAGAGCA-3’, and reverse, 5’-TAGCATCTCGGCTGGACTTCGA-3’.

### Organoid dissociation for single cell RNA-seq (scRNA-seq)

Cultured organoids were harvested after 24 hrs post-infection and then washed in cold PBS to remove Matrigel. The organoids were incubated with pre-warmed TrypLE (Thermo Fisher Scientific) for 30 min in a water bath at 37°C with consistent microscopic examination. When cells reached a single-cell state, they were resuspended in DMEM/F12 and spun at 500 g for 5 min at RT. The supernatant was removed by the pipet, and the cell suspensions were resuspended in PBS containing 0.04% BSA. The suspensions were passed through a 30 μm cell strainer and then washed with PBS containing 0.04% BSA 3 times. Resulting cell suspensions were used for a single-cell RNA sequencing kit from 10xGenomics (Pleasanton, CA, USA). A scheme of the single cell collection and scRNA-seq is described in Figure S1.

### scRNA-seq library preparation

Single-cell suspensions were loaded onto the 10x Chromium controller using the 10xGeomics Single Cell 3’ Library Kit v.2 (10xGenomics) and Chromium Next GEM Single Cell 3’ Reagent Kits v.3.1 (10xGenomics) following the manufacturer’s guideline. Briefly, single cells and beads were generated, followed by reverse transcription, cDNA amplification, fragmentation, and ligation with adapters with unique sample index PCR. Constructed libraries were sequenced with Illumina NovaSeq by Novogene Company Limited (Cambridge, UK).

### scRNA-seq data processing

Seurat 4.3.2 was used for clustering. After filtering out cells with gene number < 500, gene number ranked in the top 1%, and a mitochondrial gene ratio of < 15%. The dimension was reduced by principal component analysis (PCA) and the data were visualized by tSNE or unsupervised cluster UMAP.

### Data format for RAW data

Single cells are loaded onto the Chromium controller (10X Genomics) to partition single cells into gel beads. Single-cell transcriptomic libraries are generated using Chromium next GEM single cell 3’ kits with dual indexes for each library (V3.1 10X Genomics) according to the manufactory’s instructions. Pre-made libraries are sent to Novogene for Illumina NovaSeq sequencing with 50 M depth reads per sample for generating Fastq files with cloud delivery up to 80 GB.

### Data format and outsource libraries for downstream analysis

10X gene expression data are first accessed using Cell Ranger (v.8.0.1, 10X Genomics) using a customized version of human reference (GRCh38) in LINUX system. Samples are aggregated and normalized by the median number of mapped reads per identified cell. Data under h5 files then are imported into Seurat (v.4.3.2) for downstream analysis and data visualization. After filtering out cells with gene number < 200, nFeature counts over 4500, and a mitochondrial gene ratio of < 15% (Figure S2). The dimension was reduced by principal component analysis (PCA) and the data were visualized by tSNE or unsupervised cluster UMAP. The resulting SCT corrected counts were used for UMAP visualization and clustering downstream analysis for quantitative comparison. principal component analysis (PCA) using 3000 height variable genes based on average gene expression. The top principal components were used to construct a shared closet neighbor (SNN) graph to obtain 6 clusters in single cell population. UMAP visualization was calculated using 30 neighboring points for the manifold structure. Cell types are annotated based on markers genes for every cell type identified by comparing the expression of each gene in a given cluster against the rest of the cell population with the supplement information of SingleR function (SingleR v 2.6.0) with outsource library from Human Gut Atlas (https://www.gutcellatlas.org/) in R environment. To determine genes that are considered as markers for cluster, Leiden clustering is performed with *P adjust* < 0.05. Reactome enrichment of cluster markers are performed with Benjamini-Hochberg multiple testing adjustments with *P adjust* less 0.05 by using the top marker genes of the clusters with the visualization using R package. Different gene expression (DEGs) of two study groups as CVBs-infected and Control-groups were calculated following Wilcox test with a threshold of log2FC more than 0.25 and pajust less than 0.05 for indicating top gene expression chance between cell population.

## RESULTS

### Interferon-Driven Antiviral Programs and TG2 Upregulation Define the Organoid Response to CVB1

Gluten-induced inflammation in CeD encompasses high levels of IFN-γ expression^22^ and IFN-γ has been shown to induce TG2 expression in intestinal epithelial cancer cells in vitro.^16^ We have also shown that TG2 expression positively correlates with the epithelial IFN-γ-response in CeD patients challenged with gluten and TG2 expression is significantly induced in human intestinal organoids treated with IFN-γ,^9^ Since there is growing evidence that type II class of interferon (IFN-γ) induces TG2 expression we asked if type I and III interferons e.g. IFN-β and IFN-λ produced in the intestinal epithelium in cis, during virus infection, is also associated with increased TG2 mRNA expression.

CVB1 infection triggered a coordinated antiviral reaction in human intestinal organoids, revealing the innate capacity of the epithelium to sense and respond to enteroviral challenge. As viral RNA accumulated, epithelial cells rapidly engaged their intrinsic defense machinery, marked by the early and robust induction of type I (IFN-β) and type III (IFN-λ2/3) interferons (Fig. 1A). This interferon surge unfolded in parallel with the rise in *TGM2* expression, highlighting that TG2 is not solely a downstream target of IFN-γ-driven inflammation but is also intimately coupled to epithelial interferon signaling during active viral infection. Moreover, canonical interferon response genes are induced following CVB1 infection (Figure S3). Gene set enrichment analysis revealed activation of metabolic pathways, including oxidative phosphorylation (NES = 3.86, FDR = 6.16 × 10^−15^) (Figure S4), consistent with the energetic demands of interferon-driven antiviral responses.^23^ These findings position TG2 as an integral component of the mucosal antiviral landscape, linking viral sensing directly to one of the key enzymes implicated in celiac disease pathology.

**Figure 1.**
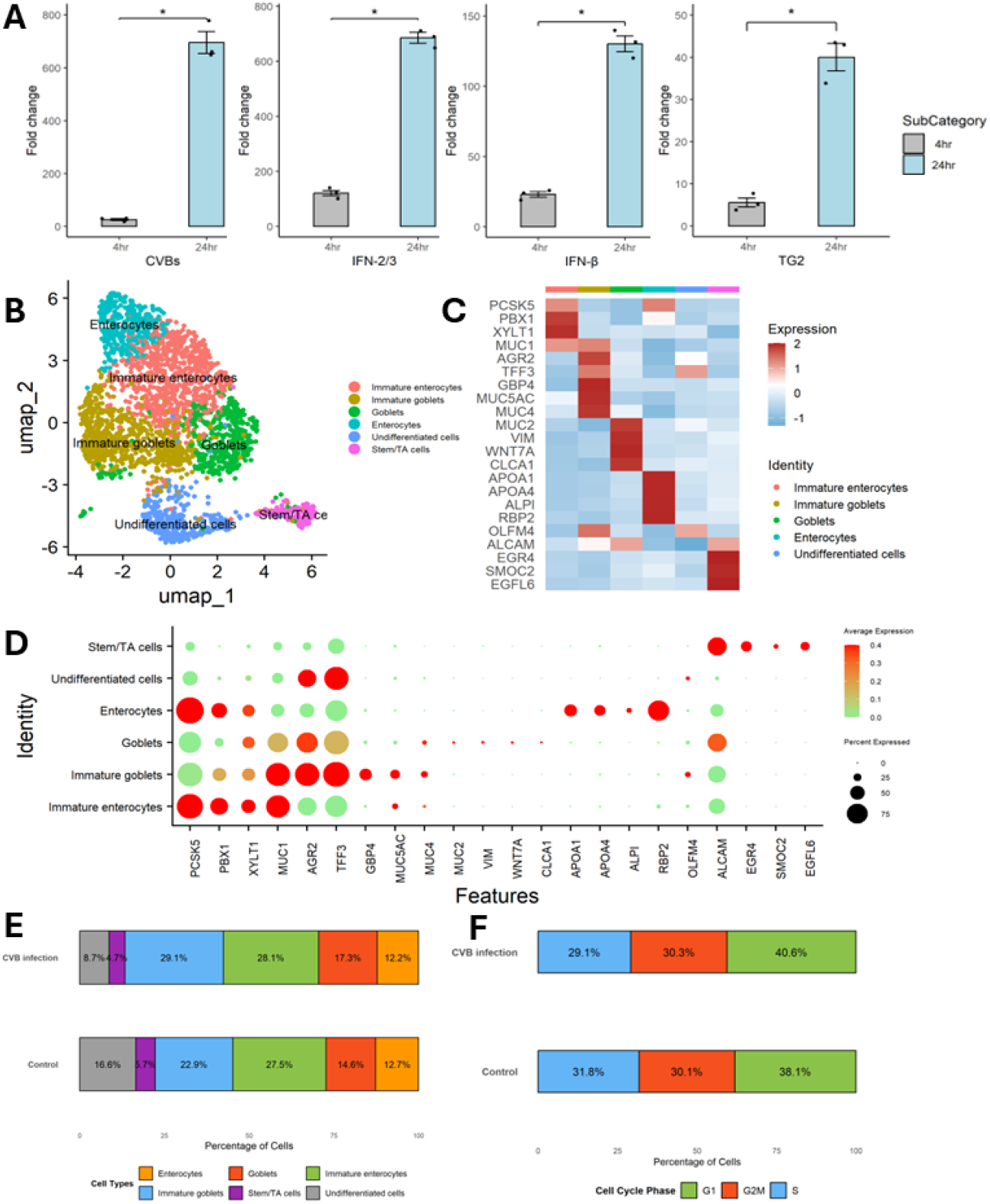
CVB1 infection induces a robust interferon response and TG2 upregulation while preserving epithelial cell diversity in human intestinal organoids. **A)** RT–qPCR quantification of viral RNA (CVB1) confirms efficient infection at 4 and 24 hours post-infection. Expression of type I (IFN-β) and type III (IFN-λ2/3) interferons, as well as *TGM2*, is significantly upregulated following infection. Data represent three biological replicates; bars indicate mean ± SD, *P* < 0.05. **B)** UMAP projection of single-cell RNA sequencing data identifies six epithelial cell populations in human intestinal organoids, including immature enterocytes, enterocytes, goblets, immature goblets, undifferentiated cells, and stem/TA cells. **C)** Heatmap showing scaled expression of representative marker genes across identified epithelial cell populations, supporting cell-type annotation. **D)** Dot plot of canonical lineage marker expression across epithelial clusters. Dot size represents the percentage of cells expressing each gene, and color intensity indicates average expression levels. **E)** Relative proportions of epithelial cell types in control and CVB1-infected organoids, demonstrating infection-associated shifts in cell-type composition without loss or emergence of major epithelial lineages. **F)** Distribution of epithelial cells across cell cycle phases (G1, S, and G2/M) in control and CVB1-infected organoids, indicating that viral infection does not markedly alter cell cycle dynamics.

To understand how this antiviral state reshapes the epithelial environment at the single-cell level, we performed scRNA-seq on infected and uninfected organoids. The analysis revealed a transcriptionally rich and diverse epithelium, composed of six major epithelial cell types that remained present across both conditions (Fig. 1B-D). CVB1 infection did not erase or create lineages but subtly reorganized the epithelial community: undifferentiated cells diminished, while immature goblet cells became more prominent, suggesting selective sensitivity or differential activation of secretory precursors during infection (Fig. 1E). Notably, these shifts occurred without major alterations in cell-cycle dynamics, indicating that CVB1 acts primarily by reprogramming epithelial identities and antiviral states rather than perturbing proliferation (Fig. 1F). Together, these data outline a cohesive narrative in which CVB1 activates a powerful interferon-driven antiviral program, induces TG2 as part of this response, and remodels epithelial composition in a manner that may have broader implications for mucosal immunity and celiac disease susceptibility.

### CVB1 targets goblet-lineage precursors and triggers interferon-driven TG2 and MUC5AC induction

We next examined how CVB1 exposure reshapes viral entry factor expression and antiviral signaling at single-cell resolution. Feature mapping on the UMAP revealed that the primary entry receptor CXADR (CAR) and the co-receptor CD55 (DAF) are not uniformly expressed but instead show lineage-skewed distributions across epithelial populations, consistent with receptor-defined susceptibility niches (Fig. 2A). To determine whether infection alters these entry landscapes, we compared *CXADR* and *CD55* between uninfected and CVB1-infected conditions. The condition-split heatmaps showed infection-associated remodeling of receptor expression patterns at the epithelial level, indicating that viral challenge is accompanied by measurable shifts in entry/co-receptor availability (Fig. 2B).

**Figure 2:**
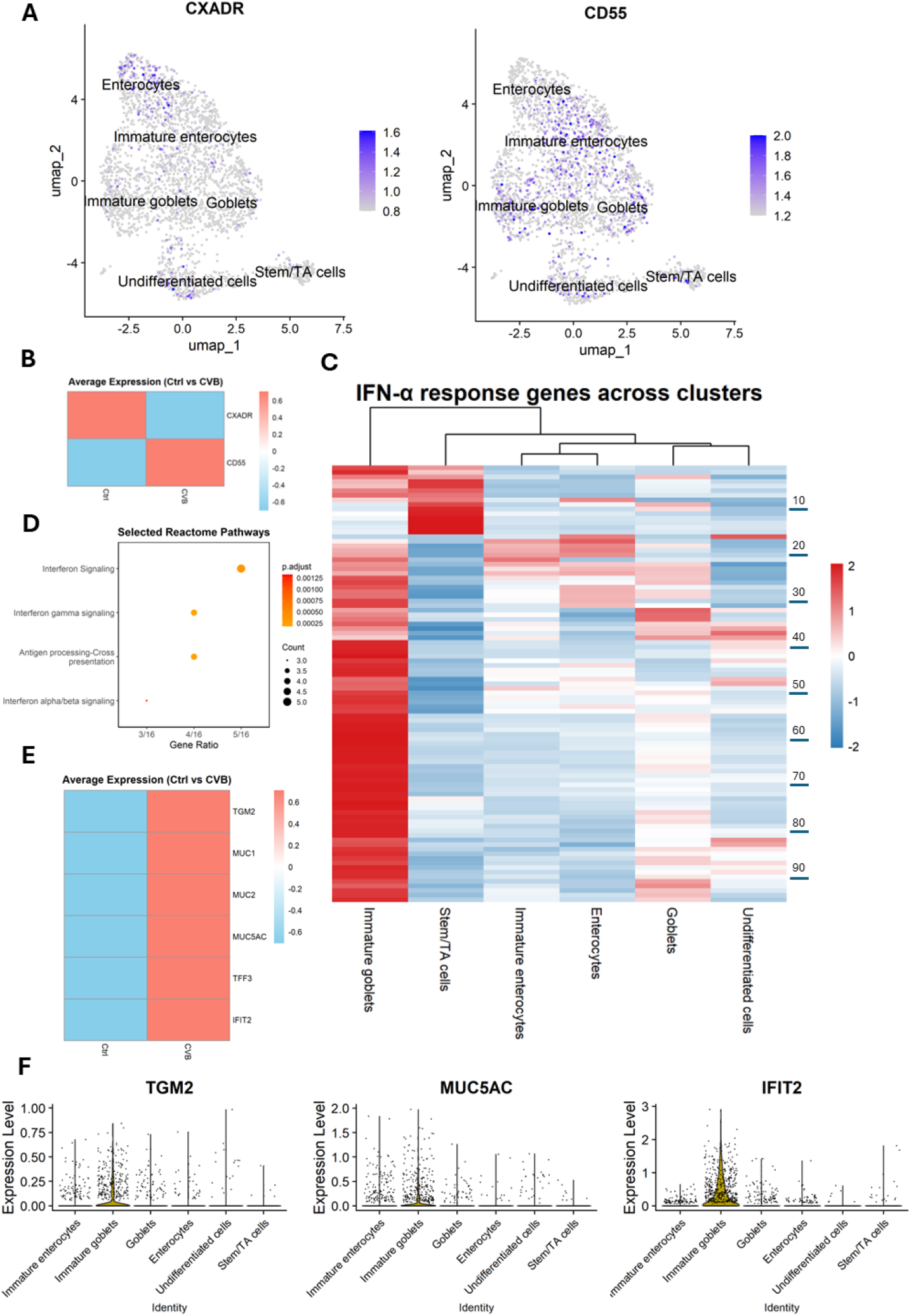
CVB1 modulates epithelial receptor expression and induces antiviral and stress-responsive transcriptional programs in goblet-lineages in human intestinal organoids. **A)** UMAP projection illustrating the distribution of the CVB1 entry receptor CXADR (CAR) and co-receptor CD55 (DAF) across epithelial cell populations in human intestinal organoids, highlighting lineage-restricted receptor enrichment. **B)** Condition-separated heatmaps showing differential expression of CXADR and CD55 in uninfected versus CVB1-infected organoids, indicating infection-associated remodeling of receptor landscapes. **C)** Interferon-stimulated gene (ISG) heatmap summarizing antiviral transcriptional activation across epithelial clusters. Immature goblet-lineage cells exhibit the strongest ISG induction following CVB1 infection. Rows correspond to proteins indexed 1– 95. Full protein names are provided in Supplementary Table S1. **D)** GO biological process enrichment of upregulated genes in infected organoids (Wilcoxon, p < 0.05), identifying antiviral pathways including type-I interferon signaling and antigen-processing modules. **E)** Condition-split heatmaps comparing TG2, mucins and IFIT2 expression between uninfected and CVB1-infected organoids, illustrating infection-associated remodeling of the immature goblet cell population. **F)** Violin plot showing the specificity of TG2, MUC5AC and IFIT2 induction in immature goblet cell population after CVB1 infection.

Next, we asked which epithelial subsets mount the strongest antiviral response. A cluster-wise interferon-stimulated gene (ISG) heatmap demonstrated pronounced ISG activation in defined epithelial lineages following CVB1 infection, highlighting immature goblet–lineage cells as prominent responders within the epithelium (Fig. 2C). At the pathway level, GO biological process enrichment of upregulated genes in infected organoids underscored antiviral programs, including type-I interferon signaling and antigen-processing modules, consistent with an epithelial-intrinsic antiviral state (Fig. 2D).

We then evaluated hallmark epithelial stress and differentiation markers together with TG2 and a sentinel ISG to connect antiviral activation with lineage remodeling. CVB1 infection is accompanied by upregulation of a coherent epithelial module, including *TGM2* (TG2), *MUC1, MUC2, MUC5AC, TFF3*, and *IFIT2* (Fig. 2E). Notably, *MUC5AC*—a gastric-type mucin not typically abundant in the small intestine—rises alongside TG2 and the ISG *IFIT2*, indicating that antiviral signaling and secretory/mucin remodeling are engaged in tandem during CVB1 challenge. Finally, to localize this response, the co-induction of *TGM2, MUC5AC*, and *IFIT2* is concentrated in the *immature goblet* population after infection (Fig. 2F). This cell-type specificity integrates the earlier observations—receptor-defined access (A–B), interferon prominence (C), and antiviral pathway activation (D)—into a single lineage-centered model, wherein immature goblets serve as a key epithelial node linking viral sensing to interferon programs, TG2 induction, and mucin-layer remodeling

Together, these data show that CVB1 acts on pre-existing epithelial heterogeneity: entry factors map to distinct lineages, immature goblets mount the strongest interferon response, and the TG2–MUC5AC–IFIT2 triad is preferentially induced in this population—providing a mechanistic bridge from viral engagement to TG2 activation and secretory differentiation changes within the intestinal epithelium

### CVB1 induces an interferon-linked gastric-type mucin shift in immature goblets without pyloric differentiation

Having localized interferon activation, TG2 induction, and MUC5AC upregulation to immature goblet cells (Fig. 2), we next asked whether CVB1 infection engages a broader gastric-type mucin or epithelial “flaring” program in this population. Previous work has identified MUC6^+^ metaplastic cells (INFLAREs) from inflamed intestines from celiac disease patients.^24^ We therefore examined the expression of mucin-associated, gastric surface, and epithelial stress markers at single-cell resolution.

CVB1 infection robustly increased the expression of the gastric surface/foveolar-type mucin *MUC5AC*, together with *TGM2* and associated markers including *TFF1, CEACAM7, DUOX2*, and *LCN2* (Fig. 3A). In contrast, canonical pyloric and neck-cell markers such as *MUC6* and *AQP5* were not induced, indicating that CVB1 does not trigger a full pyloric lineage program (data not shown). To integrate these transcriptional changes at the pathway level, we performed gene-set enrichment analysis using an epithelial interferon-linked flaring (INFLARE) signature.^24^ CVB1 infection resulted in strong enrichment of the INFLARE program (NES = 3.11, FDR = 7.06 × 10^−8^; Fig. 3B). Per-cell INFLARE scores were significantly higher in infected organoids compared with controls (Wilcoxon BH *p. adj*. =1.54 × 10−^31^; Fig. 3C) and were highest within the immature goblet population (Fig. 3D). Importantly, expression of the intestinal lineage transcription factor CDX2 was not reduced upon CVB1 infection, indicating that gastric-type mucin induction occurs without loss of intestinal epithelial identity (Data now shown).

**Figure 3.**
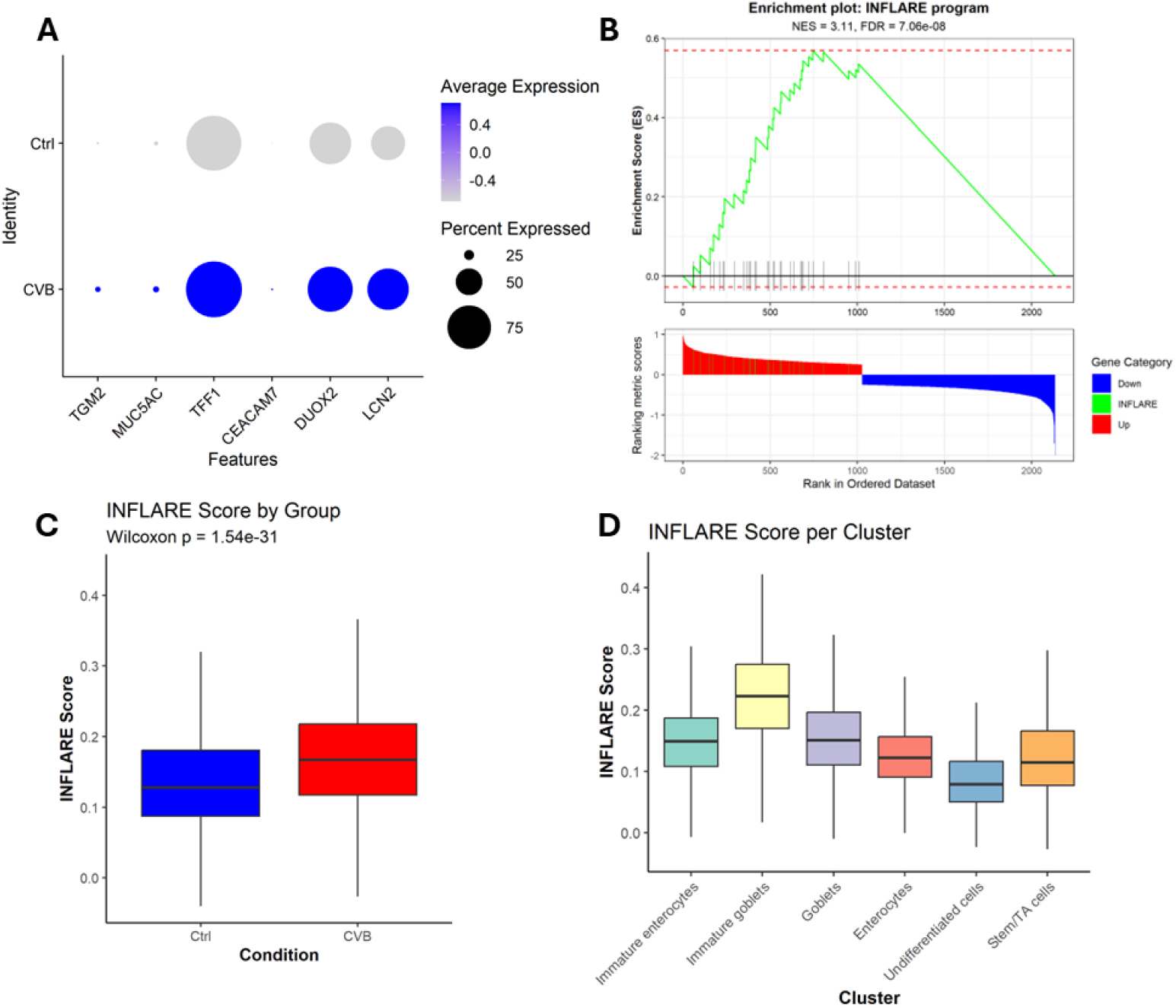
CVB1 induces an interferon-linked gastric surface/foveolar-type mucin program (INFLARE) centered on immature goblets, without pyloric/neck-cell markers or CDX2 loss. **A)** Dot plots show increased TGM2, MUC5AC, TFF1, CEACAM7, DUOX2, and LCN2 in CVB1-infected vs. control organoids. **B)** GSEA shows significant INFLARE enrichment (NES = 3.11; FDR = 7.06 × 10−^8^). **C)** Per-cell INFLARE scores are higher in CVB vs. control (Wilcoxon *p* = 1.54 × 10−^31^). **D)** INFLARE scores by epithelial cluster show the highest activation in immature goblets.

Together, these data show that CVB1 induces a coherent interferon-linked gastric surface–type mucin program centered on immature goblet cells. This response is characterized by coordinated induction of TG2 together with MUC5AC and epithelial stress–associated markers, preservation of CDX2 expression, and absence of full pyloric or neck-cell differentiation, positioning immature goblets as a key epithelial node where antiviral signaling converges with selective gastric-type mucin remodeling.

### Type-I interferons induce TG2 via JAK–STAT signaling in intestinal epithelial cells

The convergence of ISG activation, TG2 induction, and a gastric-type mucin shift in immature goblets (Figs. 2–3) prompted us to test whether type-I interferons are sufficient to drive epithelial TG2 via JAK–STAT, using IFNα/β stimulations and JAK inhibition. Treatment of organoids with IFNα markedly increased STAT1 phosphorylation, while total STAT1 protein remained constant (Fig. 4A). Tofacitinib and deucravacitinib effectively blocked IFNα-induced pSTAT1, confirming reliance on canonical JAK–STAT signaling (Fig. 4A–B). We next assessed whether this signaling was sufficient to drive TG2 expression. IFNα strongly induced *TGM2* mRNA, while co-treatment with the above-mentioned JAK inhibitors prevented this increase (Fig. 4C), demonstrating that *TGM2* transcription requires JAK–STAT activation. At the functional level, type-I interferons (IFNα and IFNβ) significantly increased TG2 enzymatic activity in human intestinal organoids, measured using the HB-2-30 fluorescent probe (Fig. 4D). This IFN-driven activation was abolished by tofacitinib or deucravacitinib (Fig. 4D-E), showing that both TG2 expression and its catalytic activity are induced by JAK-dependent pathway.

**Figure 4.**
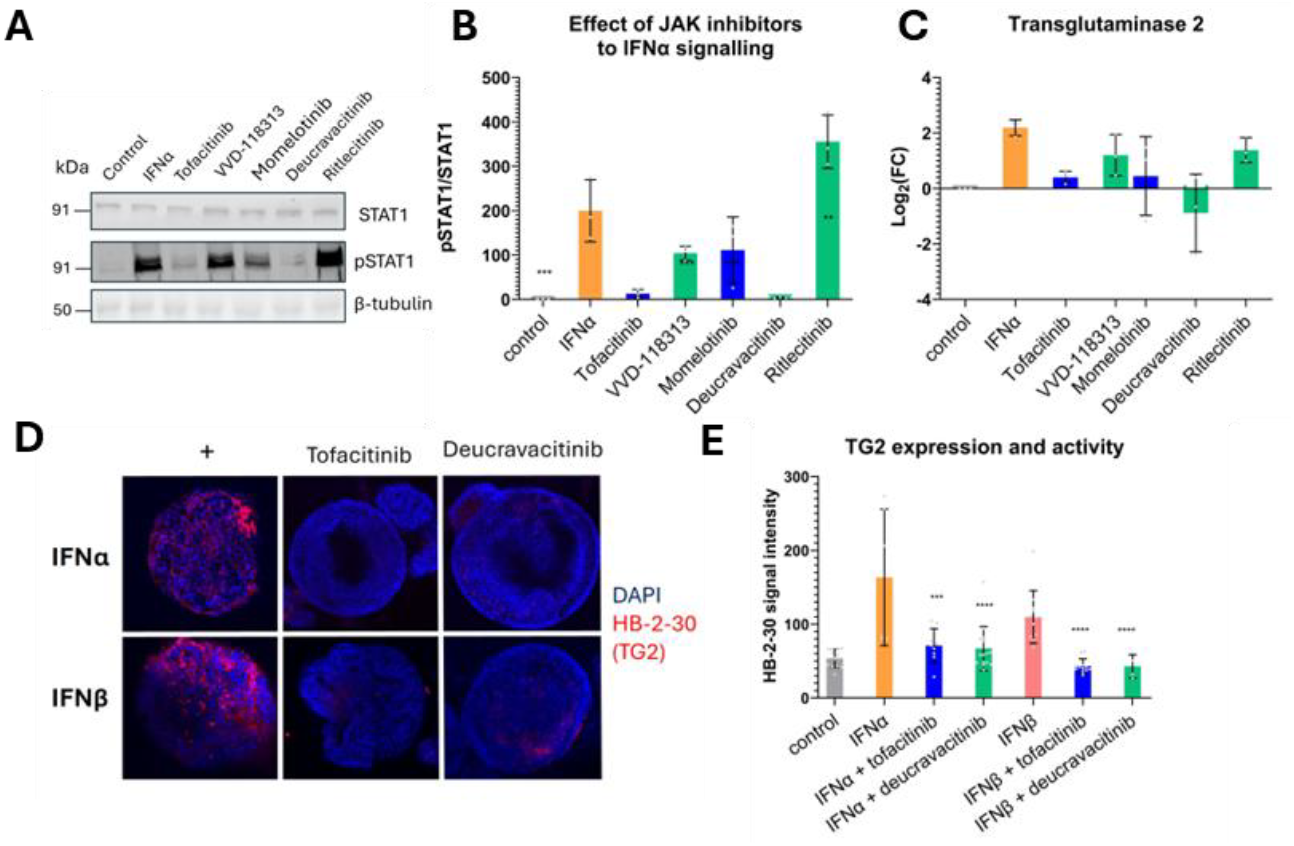
Type-I interferons induce TG2 via JAK–STAT signaling, and JAK inhibition suppresses both TG2 expression and TG2 enzymatic activity in intestinal organoids. **A)** Western blot analysis showing total STAT1, phosphorylated STAT1 (pSTAT1), and β-tubulin in organoids treated with IFNα alone or in combination with JAK inhibitors (tofacitinib, VVD-118313, momelotinib, deucravacitinib, ritlecitinib). IFNα stimulation strongly increases pSTAT1, while multiple JAK inhibitors reduce or abolish this phosphorylation. **B)** Quantification of pSTAT1/STAT1 ratios demonstrates effective suppression of IFNα-induced STAT1 phosphorylation by JAK inhibition. **C)** Relative *TGM2* mRNA levels (log_2_ fold change) in organoids treated with IFNα alone or with JAK inhibitors show that IFNα robustly induces TG2 transcription, whereas JAK inhibition blocks this induction. **D)** Confocal imaging of organoids stained with DAPI and the TG2-activity probe HB-2-30 shows increased TG2 activity following IFNα or IFNβ stimulation and marked reduction when co-treated with tofacitinib or deucravacitinib. **E)** Quantification of HB-2-30 fluorescence intensity confirms that type-I interferons increase TG2 activity and that this increase is suppressed by JAK inhibition. Error bars represent standard deviation. Statistical tests follow ANOVA/Dunnett’s post-hoc comparisons.

Together, these results demonstrate that type-I interferons are sufficient to induce TG2 via JAK–STAT, establishing a direct mechanistic link between epithelial interferon signaling and TG2 activation—precisely the pathway engaged during CVB1 infection. This provides functional evidence connecting the interferon-driven TG2 enzymatic induction, a key pathogenic step relevant to celiac disease susceptibility.

### Epithelial Cell–cell communication remains stable during interferon-driven TG2 induction

Given the strong interferon response and TG2 activation induced by CVB infection (Figs. 2– 4), we asked whether these changes were accompanied by altered epithelial cell–cell communication. CellChat analysis^25^ showed that epithelial communication networks were largely preserved, with comparable interaction numbers and strengths between control and CVB-infected organoids (Fig. 5A–B). The same epithelial populations formed the dominant interaction hubs in both conditions, indicating that communication structure is maintained despite infection. Pathway-level analyses revealed largely shared signaling pathways with only minor redistribution among low-information-flow pathways (Figure S5), and no emergence of interferon- or virus-specific ligand–receptor signaling. These data indicate that interferon-driven TG2 induction occurs primarily through cell-intrinsic mechanisms rather than through rewired epithelial communication networks.

**Figure 5.**
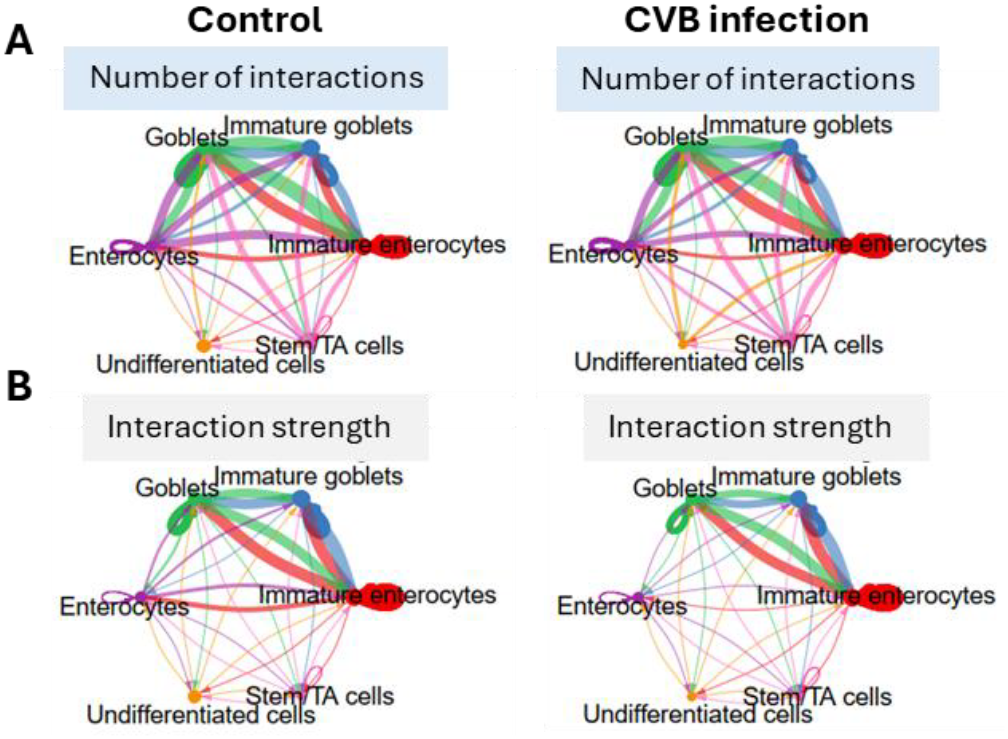
Epithelial cell–cell communication is largely preserved during CVB infection. Circle plots showing the number of interactions (**A**) and interaction strength (**B**) among epithelial cell populations in control (left) and CVB-infected (right) organoids. Overall network topology and dominant interaction hubs are similar between conditions.

## DISCUSSION

Epidemiological studies have consistently linked enterovirus infections to an increased risk of celiac disease (CeD), ^12–14,26,27^ yet the molecular basis of this association has remained unclear. In this study, we used human intestinal organoids and single-cell transcriptomics to define an epithelial-intrinsic mechanism by which coxsackievirus B1 (CVB1) infection may contribute to CeD pathogenesis. Our findings demonstrate that enterovirus infection activates a type I interferon–driven antiviral program that induces transglutaminase 2 (TG2) expression and enzymatic activity via JAK–STAT signaling, thereby linking viral sensing directly to a central biochemical step in CeD.

A key advance of this work is the identification of TG2 regulation as part of an intestinal epithelial antiviral response. While previous studies have shown that interferon-γ can induce TG2 expression, our data demonstrate that type I interferons produced locally in the intestinal epithelium are sufficient to drive TG2 expression and activity through canonical JAK–STAT signaling. This establishes TG2 as a downstream component of epithelial interferon responses, expanding its role beyond inflammation-associated contexts into early antiviral defense. Consistent with this broader role, viral infections can engage TG2 and reshape epithelial– immune interactions. RSV induces activation and release of TG2 in airway epithelium,^28^ whereas enteric viruses such as reovirus disrupt oral tolerance to dietary antigens via interferon-dependent mechanisms.^29^ Together, these findings support a general role for viral infection in coordinating TG2 activity and immune responses to luminal antigens. Importantly, we show that pharmacologic inhibition of JAK signaling effectively suppressed TG2 induction, highlighting the pathway’s specificity and potential therapeutic relevance.

Our single-cell analyses further reveal that TG2 induction is not uniform across the epithelium but is concentrated in immature goblet-lineage cells, which mount the most robust interferon-stimulated gene response following CVB1 infection. This lineage-specific response suggests that epithelial heterogeneity critically shapes how antiviral signaling is translated into functionally relevant responses. Goblet cells are central to mucosal barrier function, and their selective engagement during viral infection may create localized microenvironments enriched in TG2 activity and modified mucins, potentially amplifying exposure of the immune system to deamidated gluten peptides (Fig. 6). In addition to their barrier function, goblet cells can mediate antigen transfer through goblet cell–associated antigen passages (GAPs), a regulated transcytotic process that facilitates delivery of luminal antigens to underlying immune cells.^30^ Although TG2 can couple to LDL receptor-related protein 1 (LRP-1)-mediated endocytic pathways to facilitate antigen uptake and presentation in professional antigen-presenting cells,^20^ the absence of LRP induction in CVB1-infected epithelium (data not shown) suggests that TG2 upregulation here reflects a distinct interferon-driven epithelial program rather than engagement of receptor-mediated uptake pathways.

**Figure 6.**
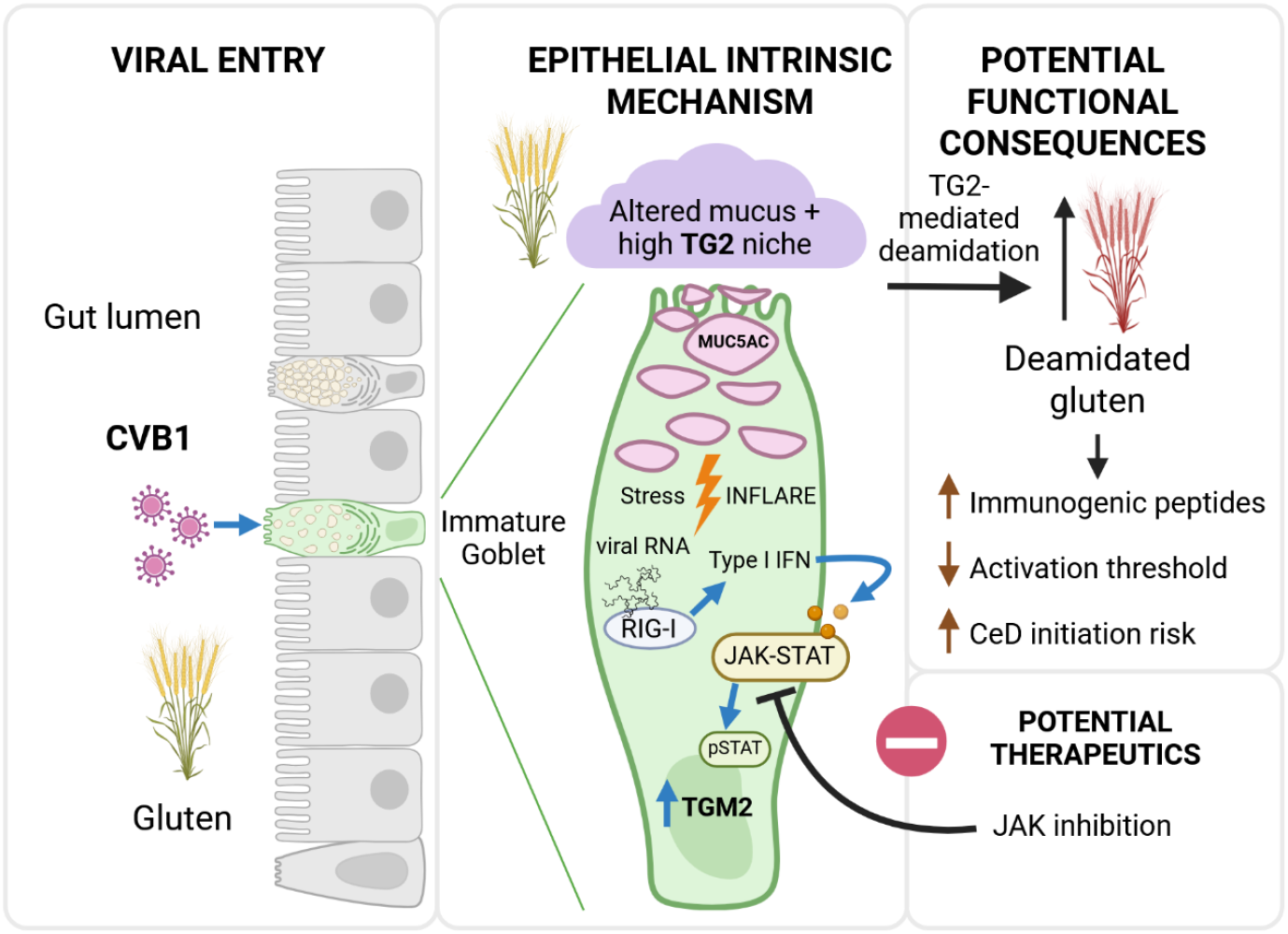
A model linking enterovirus-induced epithelial interferon signaling to TG2 activation and celiac disease priming. CVB1 infection induces a type I interferon–driven epithelial response that activates TG2 via JAK–STAT signaling, particularly in immature goblet cells. This response is coupled to mucin remodeling and creates a TG2-high epithelial niche that enhances gluten deamidation, increasing immunogenic peptide generation and lowering the threshold for T cell activation, thereby promoting celiac disease initiation. JAK inhibition blocks this pathway.

In parallel with TG2 induction, CVB1 infection triggered a coordinated epithelial remodeling program characterized by upregulation of gastric-type mucins, including MUC5AC, and enrichment of interferon-linked epithelial stress signatures. This response resembles recently described inflammatory “flaring” programs in the intestine,^24^ and is consistent with epithelial changes observed in celiac disease, where gastric metaplasia and MUC5AC-expressing cells emerge in the duodenal mucosa.^31^ The co-occurrence of TG2 activation and mucin remodeling within the same epithelial subpopulation supports a model in which antiviral responses simultaneously modify both enzymatic processing and barrier properties, two factors that are critical in shaping immune exposure to dietary antigens.

These findings provide a mechanistic framework linking enterovirus infection to CeD pathogenesis. Enteroviruses replicate within the intestinal mucosa and can induce sustained interferon responses.^32,33^ Our data suggest that, during such infections, epithelial interferon signaling directly increases TG2 activity, thereby potentially increasing the local capacity for gluten peptide modification. In the context of genetic susceptibility and dietary gluten exposure, this could lower the threshold for activation of gluten-specific T cells and promote the initiation of CeD-associated immune responses (Fig. 6). This model is consistent with prospective studies indicating that the timing of enterovirus infection relative to gluten exposure influences disease risk.^11,13^

An important implication of our findings is that TG2 induction in this context occurs independently of immune cell involvement, emphasizing the active role of the epithelium in initiating disease-relevant processes. This epithelial-intrinsic mechanism provides a potential explanation for how environmental triggers can act at early stages of disease development before overt inflammation is established. Notably, the reversibility of TG2 induction by JAK inhibition highlights epithelial interferon signaling as a tractable therapeutic target. While JAK inhibitors are widely used in other inflammatory diseases,^34^ their application in celiac disease has remains underexplored, particularly in epithelial contexts and early disease mechanisms. Our findings therefore support the repurposing of JAK inhibitors as a strategy to prevent or dampen virus-induced priming of celiac disease by directly targeting epithelial stress responses.

This study has limitations. Our experiments were performed using organoids derived from non-CeD donors and focused on a single enterovirus strain. Future studies should assess whether similar mechanisms operate across diverse enteroviruses and in genetically susceptible individuals. In addition, while our data demonstrate induction of TG2 and its enzymatic activity, further work is needed to directly link these changes to enhanced gliadin deamidation and subsequent immune activation. Incorporating immune components into organoid systems or validating findings in human biopsy material will be important next steps.

In conclusion, we identify a direct epithelial pathway linking enterovirus infection to TG2 activation via interferon-driven JAK–STAT signaling. This mechanism provides a biologically plausible bridge between viral exposure and a key initiating step in CeD pathogenesis and highlights epithelial antiviral signaling as a potential target for preventive or therapeutic intervention.

## Supporting information

list of HEDIMED investigators

## AUTHOR CONTRIBUTIONS

KV and HHL conceptualized the study. KV and HHL drafted the manuscript. HHL, VR, RD, VD, LMZ, ASK, AV and KV performed investigation, data analysis and figure generation.

JEL, CK, OS, HH and KV provided resources and data curation. KV and HH acquired the funding. All authors read and approved the final paper

## DATA AND CODE AVAILABILITY

All data are available in the main text or the supplementary materials.

## ETHICS APPROVAL

Human duodenal tissues for establishing organoid cultures used in this study were sourced from deidentified surgical specimens of the duodenum obtained from participants who had undergone biopsy procedures unrelated to CeD at Tampere University Hospital. The protocol was approved by the Ethics Committee of Tampere University Hospital (ETL code R18082).

## ACKNOWLEDGEMENTS

This project has received funding from the European Union’s Horizon 2020 Research and Innovation Programme HEDIMED project under Grant Agreement 874864 and Horizon Europe Research and Innovation Programme ENT1DEP project under Grant Agreement 101137457. Work was also supported by the Research Council of Finland (Grant no. 370828, 335437), University of Oulu and Research Council of Finland Profi8 (Grant no. 365202), the National Institutes of Health (R01 DK063158) the Finnish Cultural Foundation, Mary och Georg C. Ehrnrooths Stiftelse, Tampere tuberculosis foundation, Sigrid Juselius foundation, Finnish cancer foundation, and the State funding for university-level health research, Tampere University Hospital, Wellbeing services county of Pirkanmaa / Project No. T63474, T66984, X50020 and X50040. The funding sources played no role in the design or execution of this study or in the analysis and interpretation of the data. Authors thank Pauliina Hiltunen and Kaija-Leena Kolho for providing the intestinal biopsies from which the organoids were derived. Authors acknowledge the Biocenter Finland (BF), Tampere Imaging Facility (TIF), Tampere Genomics Facility, Tampere Adult Stem Cell Organoids Facility and Tampere Virus Production Facility.

