## Supplementary material for "Enterovirus-driven interferon signaling induces epithelial TG2 via JAK-STAT: Implications for the onset of celiac disease": list of HEDIMED investigators

HEDIMED investigator group

Baylor College of Medicine, Department of Molecular Virology and Microbiology:
Hoffman Kristi, Javornik Cregeen Sara, Petrosino Joseph, Santiago Rodriguez Tasha

Charles University, 2nd Faculty of Medicine:
Cinek Ondrej, Fronkova Eva

COPSAC:
Bisgaard Hans, Bønnelykke Klaus, Brandt Sarah, Sevelsted Astrid, Shah Shiraz, Stokholm Jakob, Thorsen Jonathan

Czech University of Life Sciences Prague, Department of Food Science:
Havlik Jaroslav, Jeníčková Eliška

Empirica:
Schmidtmann Daniel, Thiel Rainer

Finnish Institute for Health and Welfare, The Department of Health Security and The Department of Public Health and Welfare:
Hakola Leena, Kiviranta Hannu, Rantakokko Panu, Virtanen Suvi M.

Gnomon:
Berler Alexander, Karabatea Apostolia, Papadopoulou Korina

Graz University of Technology, Institute of Environmental Biotechnology:
Berg Gabriele, Wicaksono Wisnu

Lund University, Department of Clinical Sciences:
Agardh Daniel, Aronsson Carin Andrén, Lundgren Markus, Michaela Boström

Natural Resources Institute Finland, Horticulture Technologies:
Roslund Marja, Sinkkonen Aki

Satellio:
Alibakhshi Sara, Häme Lauri

Swiss Center for Electronics and Microtechnology (CSEM):
Boia Patricia-Daiana, Burr Loïc, Cattaneo Stefano, Cristofollini Peter, Demuru Silvia, Generelli Silvia, Hermann Nicola, Chai-Gao Hui, Paoletti Samantha, Petkus Bradley, Ruth Edith, Shynkarenko Yevhen

Tampere University, Faculty of Medicine and Health Technology:
Antola Erika, Eurén Anna, Halonen Miia, Hyöty Heikki, Jouppila Niila, Kangasmäki Nanna, Kivelä Laura, Kurppa Kalle, Laiho Jutta, Laitinen Olli, Lehtonen Jussi, Lin Jake, Lindfors Katri, Lönnrot Maria, Malkamäki Johannes, Mäkinen Jani, Numminen Henna, Nurminen Noora, Nykter Matti, Oikarinen Sami, Oikarinen Maarit, Palmu Tiina, Puustinen Leena, Seppälä Erika, Sioofy-Khojine Amirbabak, Turppa Minna, Viiri Keijo

Tampere University Hospital:
Knip Mikael

Tartu University Hospital:
Peet Aleksandr, Simre Kärt, Tillmann Vallo

The Norwegian Institute of Public Health:
Hård af Segerstad Elin, Lund-Blix Nicolai A., Magnus Maria, Rantala Aino-Kaisa, Stene Lars, Størdal Ketil, Tapia German

University of Helsinki, Faculty of Biological and Environmental sciences:
Mäkelä Iida, Romantschuk Martin, Soininen Laura

University of Oulu, Department of Paediatrics:
Honkanen Tiia, Pietilä Juho, Rönkä Nelli, Valtanen Toni, Veijola Riitta

University of Siena, Department of Medicine, Surgery and Neuroscience:
Bargagli Elena, Dotta Francesco, Nigi Laura, Sebastiani Guido

University of Tartu, Department of Immunology, Institute of Pharmacy:
Aints Alar, Alnek Kristi, Bärenson Anu, Kirss Anne, Laidmäe Ivo, Oras Astrid, Tagoma Aili, Uibo Raivo, Vorobjova Tamara

University of Turku, Turku Bioscience Centre (University of Turku and Åbo Akademi University), Institute of Biomedicine, Department of Paediatrics:
Bojovic Iltana, Elo Laura, Hirvonen Karoliina, Junttila Sini, Kettunen Jalmari, Lahesmaa Riitta, Lempainen Johanna, Moulder Robert, Rasool Omid, Starskaia Inna, Suomi Tomi, Toppari Jorma, Ullah Ubaid

VTT Technical Research Centre of Finland Ltd:
Emilia Barannik, Gbodjo Yawogan Jean Eudes, Molinier Matthieu, Nevanen Tarja, Pajula Juha, Pärkkä Juha, Ranta Jukka, Rökman Jyri, Saviranta Petri, Ylén Peter, Parmes Eija
