## Supplemental figures S1-S5 for "Enterovirus-driven interferon signaling induces epithelial TG2 via JAK-STAT: Implications for the onset of celiac disease"

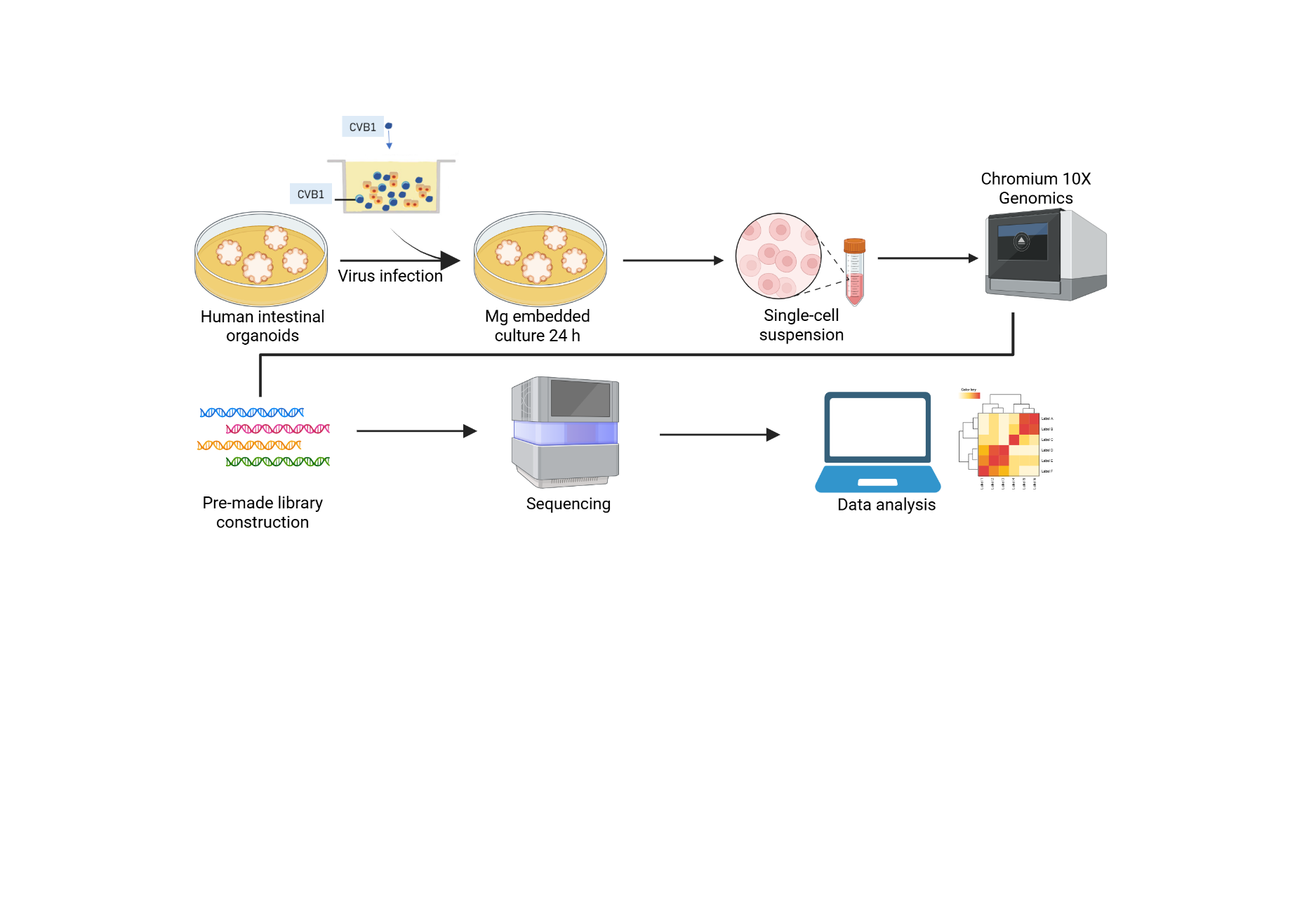

**Figure S1: Schematic presentation of the study.** Organoids collected from Matrigel were suspended with CVB1 virus following MOI 1:6 for 1 hr at RT and then embedded again in the culture environment. After 24 hrs, organoids were broken with TrypLE for collecting single cell suspension. The population was loaded to Chromium 10X Genomics as the manufacturing protocol for making library construction, which was sequenced with NovaSeq with 50M depth reads per cell. Created with Biorender.

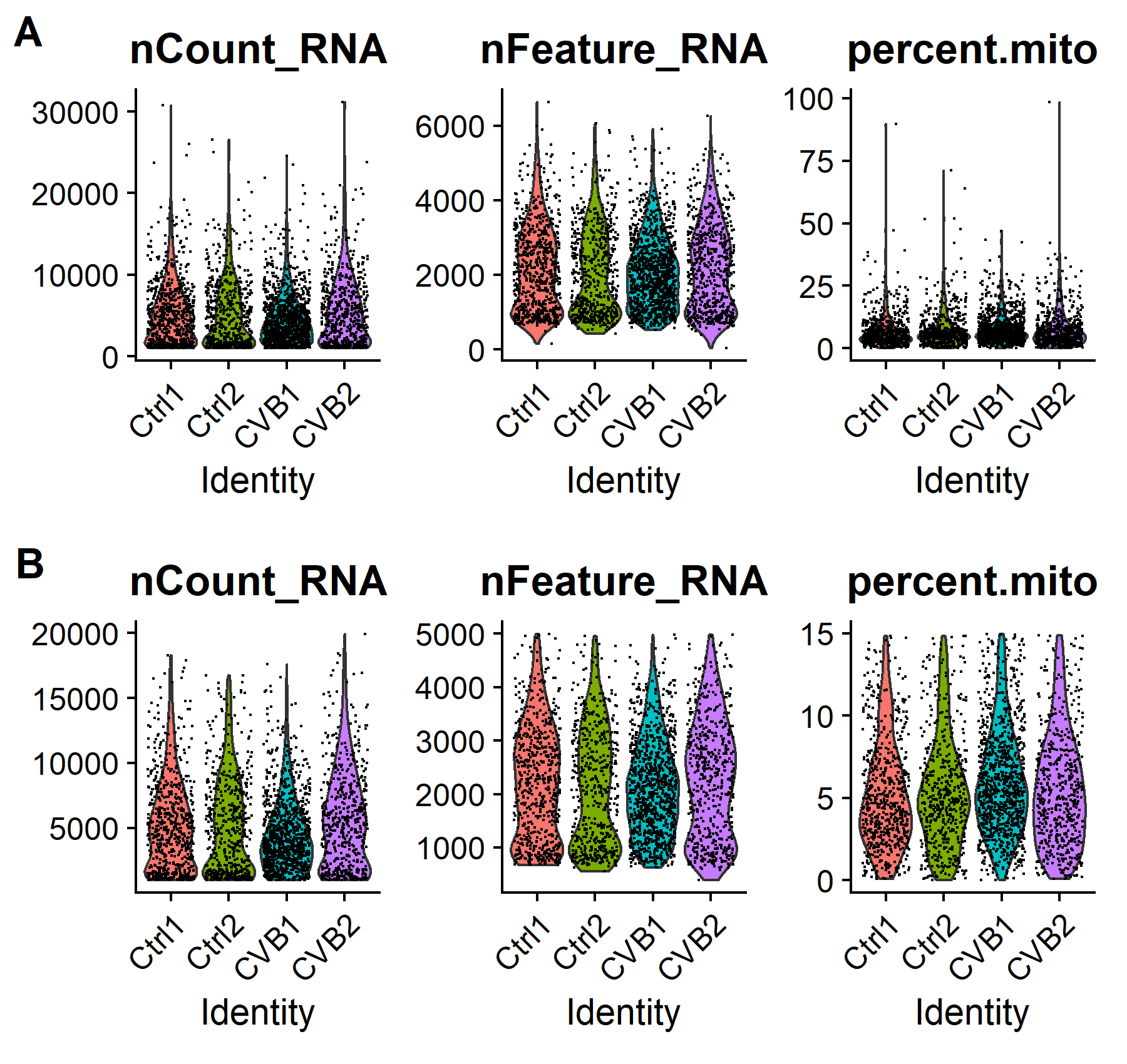

**Figure S2**: Filtering for cell quality. Total counts, nFeatures, and mitochondrial gene percentages are shown for samples before and after filtering; prefiltering (A) and postfiltering (B).

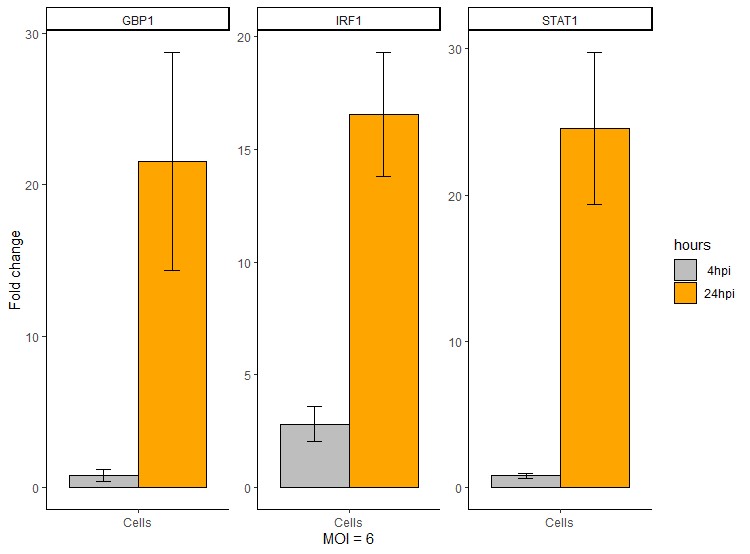

**Figure S3: Canonical interferon response genes in human intestinal organoids.**

Human intestinal organoids were infected with CVB1 and collected for mRNA extraction. The expression levels of GBP1, IRF1, and STAT1 were determined by qPCR and calculated based on the ratio between the time points with a comparison with uninfected group.

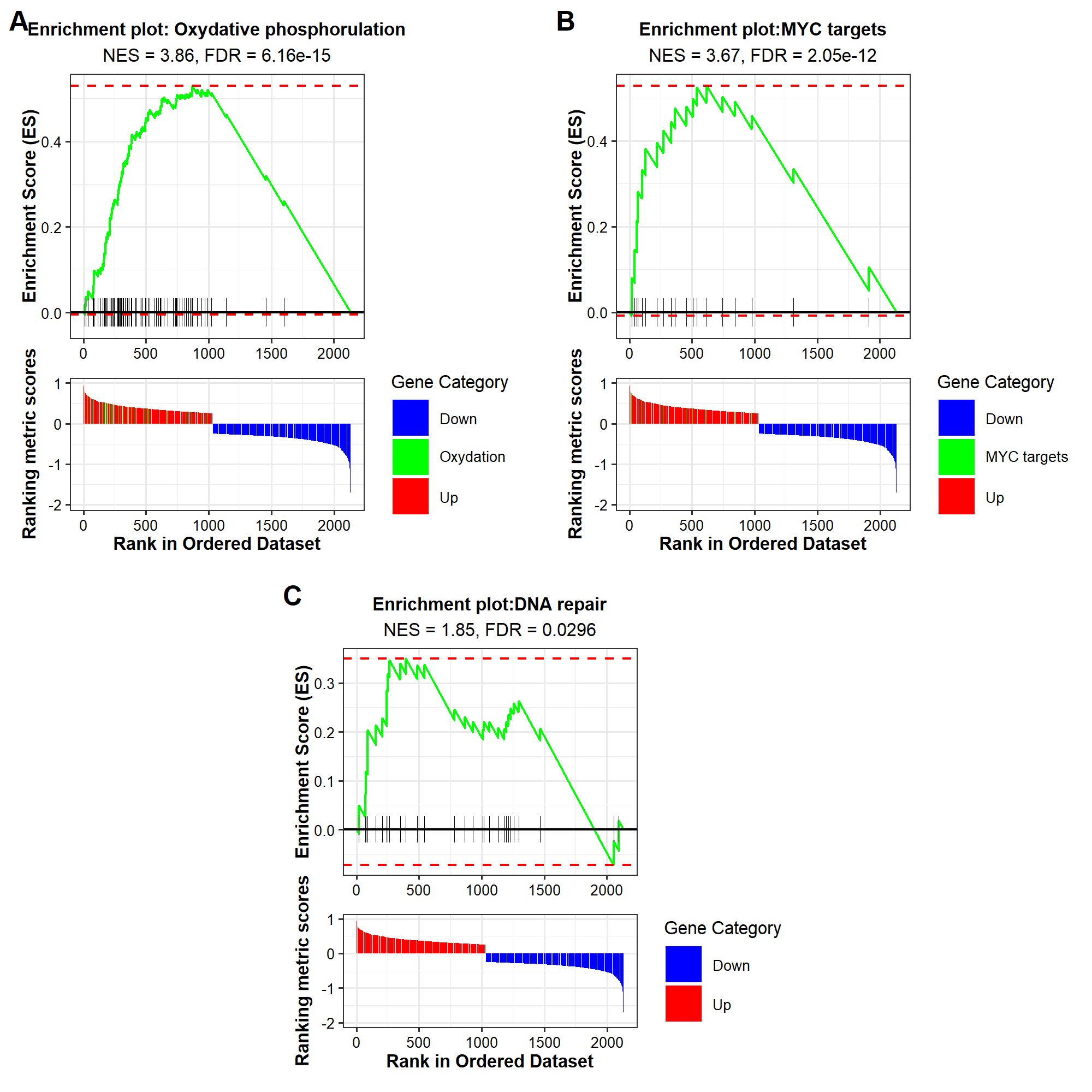

**Figure S4: Oxydative phosphorylation Reactome enrichment under CVB infection**

GSEA plots showing the enrichment scores of enriched gene sets in Oxydative phosphorylation-

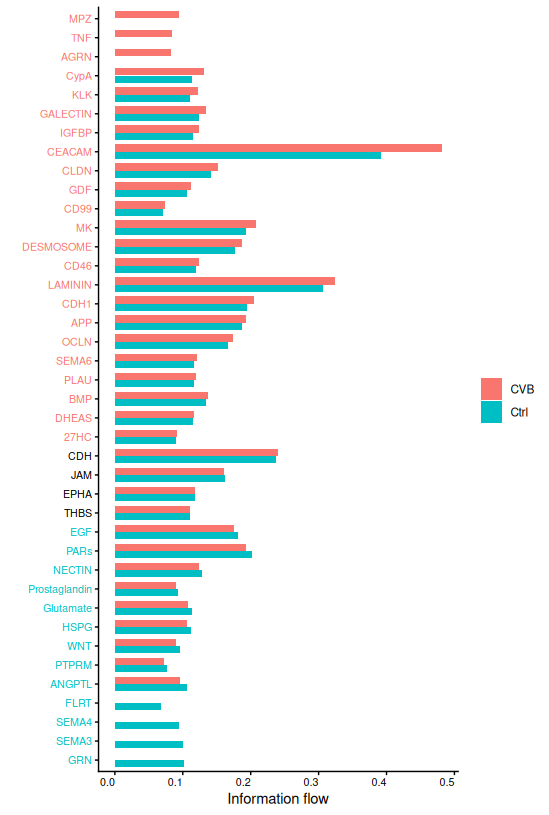

**Figure S5. Cell–cell communication analysis of epithelial organoids.** Bar plots show pathway information flow in control (teal) and CVB1-infected (red) samples. Overall signaling remains largely preserved, with modest changes in selected pathways, supporting predominantly cell-intrinsic respon
